# Passive Cell-Flux Partitioning Enables Single-Cell-Constrained Impedance Sensing From High-Density Microbial Cultures

**DOI:** 10.64898/2026.09.30.755765

**Authors:** Woud Alsadoun, Luca Pungan, Yash H. Patel, Juan J. Robayo Yepes, Yueying Yang, Rabia T. Yazicigil, Miguel Jimenez

## Abstract

Fermentation is a process through which microorganisms are used to transform basic molecules into high value substances in various applications such as biopharma, food production, and biofuel generation. In such processes, monitoring the health of microbial cells is essential, as it is a key factor in the final product yield. Traditional methods of assessing the health of fermenter populations have involved manually extracting samples from the fermenter for benchtop testing, which lacked the accuracy and efficacy of real-time continuous measurements. This has opened the door for inline sensors to be integrated into fermenters for continuous measurement; including microfluidic based Electrochemical Impedance Spectroscopy (EIS). EIS utilizes a range of frequencies to gain insight into the characteristics of microbial populations, such as cell size and membrane capacitance. EIS cytometry, in particular, has emerged as a relevant tool, as it enables single-cell level detection in real-time. However, current EIS cytometry devices cannot be deployed directly within fermenters due to the high cell density conditions. This has limited EIS analysis in fermenters to bulk measurements averaged across a population. Here, we present a microfluidic chip that uses passive cell-flux-partitioning geometry to direct cells from high-density culture through a single-cell-constrained flow path containing integrated impedance-sensing electrodes. The proposed chip combines geometric channel design and embedded electrodes to enable continuous impedance measurements from cell-containing flow while operating at input cell densities on the scale of 10^9^ CFU/mL. This work demonstrates a first step toward integrating single-cell-constrained microfluidic handling with impedance sensing for inline monitoring in industrial biomanufacturing.

## 1. Introduction

Fermentation is the 10,000 year old practice of using microorganisms to produce valuable products and plays a pivotal role in modern biomanufacturing ^1,2^. Expanding beyond agricultural processes, fermentation at an industrial level now includes medicines, materials, and fuels ^3^. These products can be easily manufactured at small scales, however, to meet global demand for bulk commodities, production must scale up significantly, with commodity fermenters exceeding 100,000 liters ^4^. This has proven to be difficult, as scaling up these benchtop reactions by several orders of magnitude is often met with complex challenges such as flow field differences, biological noise, and microbial kinetics variations within the bioreactor ^5–7^. Therefore, maintaining performance at industrial scales requires continuous in-line monitoring of fermenter conditions to achieve the intended performance and product quality ^8^. Despite the importance of continuous in-line measurements, the current parameters tracked, by and large, continue to be limited to physical measurements of the fermenter medium (temperature, pH, dissolved oxygen) ^9^. However, these methods are indirect measures of microbial health, and therefore do not capture the true cell state or account for cell-to-cell heterogeneity which arises during industrial scale fermentation ^10^. Outside of industrial fermentation, advances in single-cell analysis have revealed that individual cells within the same environment can differ significantly in morphology, gene expression, and metabolic activity among other critical characteristics ^10^. While it is clear that substantial cell-to-cell heterogeneity is present in industrial scale fermenters, it continues to be challenging to adapt single-cell methods to the high cell density environment required for in-line monitoring.

This value of single-cell monitoring in fermenters has been recognized and is evident in the development of single-cell monitoring technologies including optical spectroscopy, absorbance based probes, *in situ* microscopy with image analysis, and electrochemical impedance spectroscopy (EIS) (Figure 1A). Such methods allow the user to probe cell populations at single-cell resolution and extract vital information regarding cell state and productivity that would otherwise have to be inferred from bulk measurements^11,12^. However, to apply these technologies within fermenters would require complex alterations within the fermenter’s structure, especially for in situ microscopy which is the closest to the needed resolution, and is being used in bioprocesses ^13^. Other common single cell sorting methods are also currently not compatible with in-line monitoring due to sample preparation requirements (Table 1). The key challenge to translating these advanced tools for microbial fermenter monitoring is how to continuously isolate single cells from a population density on the scale of 10^9^ cell/mL, as typically found in fermenters ^14^.

**Table 1.**
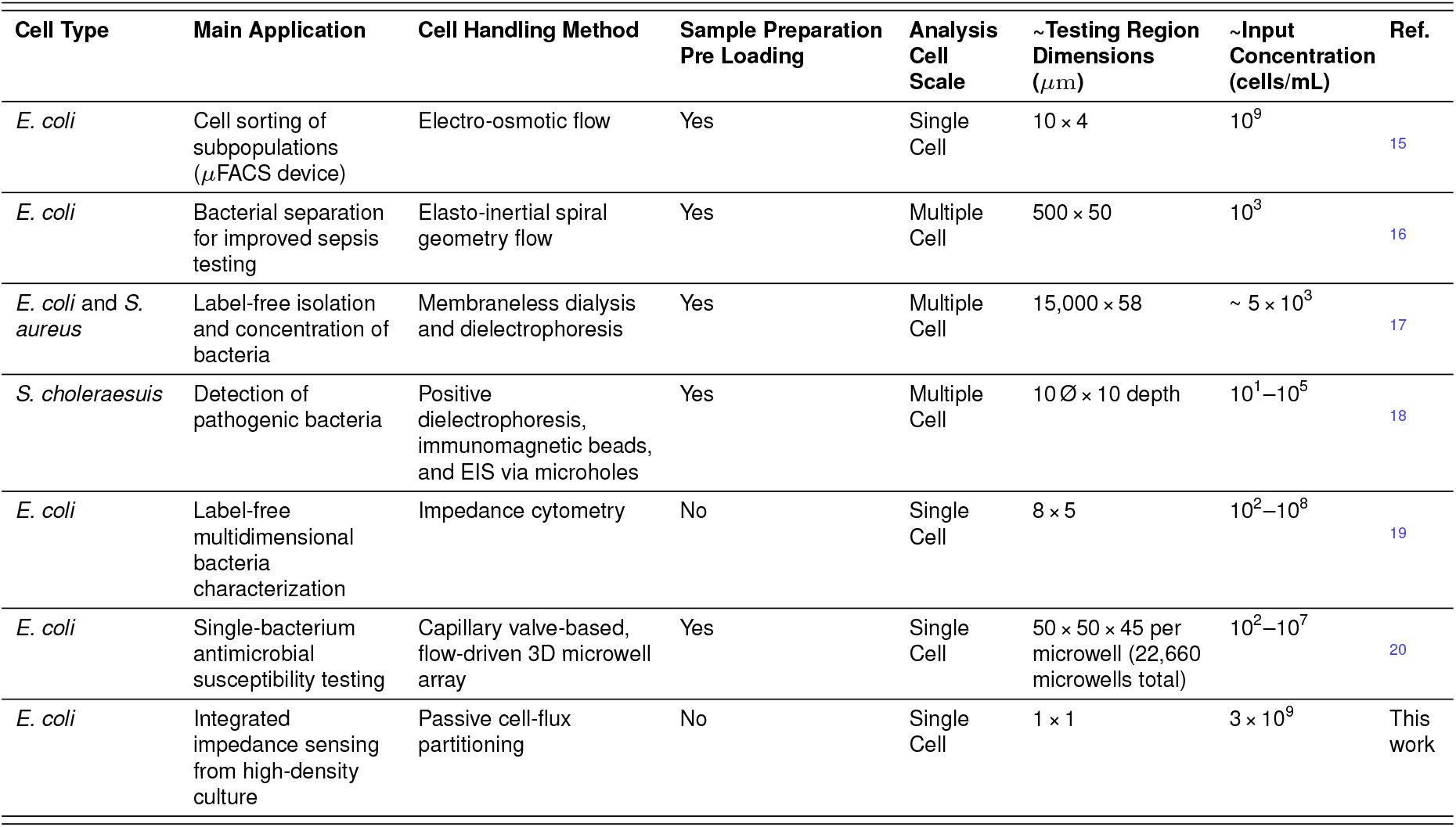
Metrics describing scale and limits of detection of varying cell separation methods described in this manuscript. Specific examples attempting to isolate cells from a high-density population are included. All methods listed are under continuous flow for comparison purposes.

| Cell Type | Main Application | Cell Handling Method | Sample Preparation Pre Loading | Analysis Cell Scale | ~Testing Region Dimensions ($\mu\text{m}$) | ~Input Concentration (cells/mL) | Ref. |
| --- | --- | --- | --- | --- | --- | --- | --- |
| *E. coli* | Cell sorting of subpopulations ($\mu\text{FACS}$ device) | Electro-osmotic flow | Yes | Single Cell | $10 \times 4$ | $10^9$ | 15 |
| *E. coli* | Bacterial separation for improved sepsis testing | Elasto-inertial spiral geometry flow | Yes | Multiple Cell | $500 \times 50$ | $10^3$ | 16 |
| *E. coli* and *S. aureus* | Label-free isolation and concentration of bacteria | Membraneless dialysis and dielectrophoresis | Yes | Multiple Cell | $15,000 \times 58$ | $\sim 5 \times 10^3$ | 17 |
| *S. choleraesuis* | Detection of pathogenic bacteria | Positive dielectrophoresis, immunomagnetic beads, and EIS via microholes | Yes | Multiple Cell | $10 \text{ } \varnothing \times 10 \text{ depth}$ | $10^1 - 10^5$ | 18 |
| *E. coli* | Label-free multidimensional bacteria characterization | Impedance cytometry | No | Single Cell | $8 \times 5$ | $10^2 - 10^8$ | 19 |
| *E. coli* | Single-bacterium antimicrobial susceptibility testing | Capillary valve-based, flow-driven 3D microwell array | Yes | Single Cell | $50 \times 50 \times 45$ per microwell (22,660 microwells total) | $10^2 - 10^7$ | 20 |
| *E. coli* | Integrated impedance sensing from high-density culture | Passive cell-flux partitioning | No | Single Cell | $1 \times 1$ | $3 \times 10^9$ | This work |

**Figure 1.**
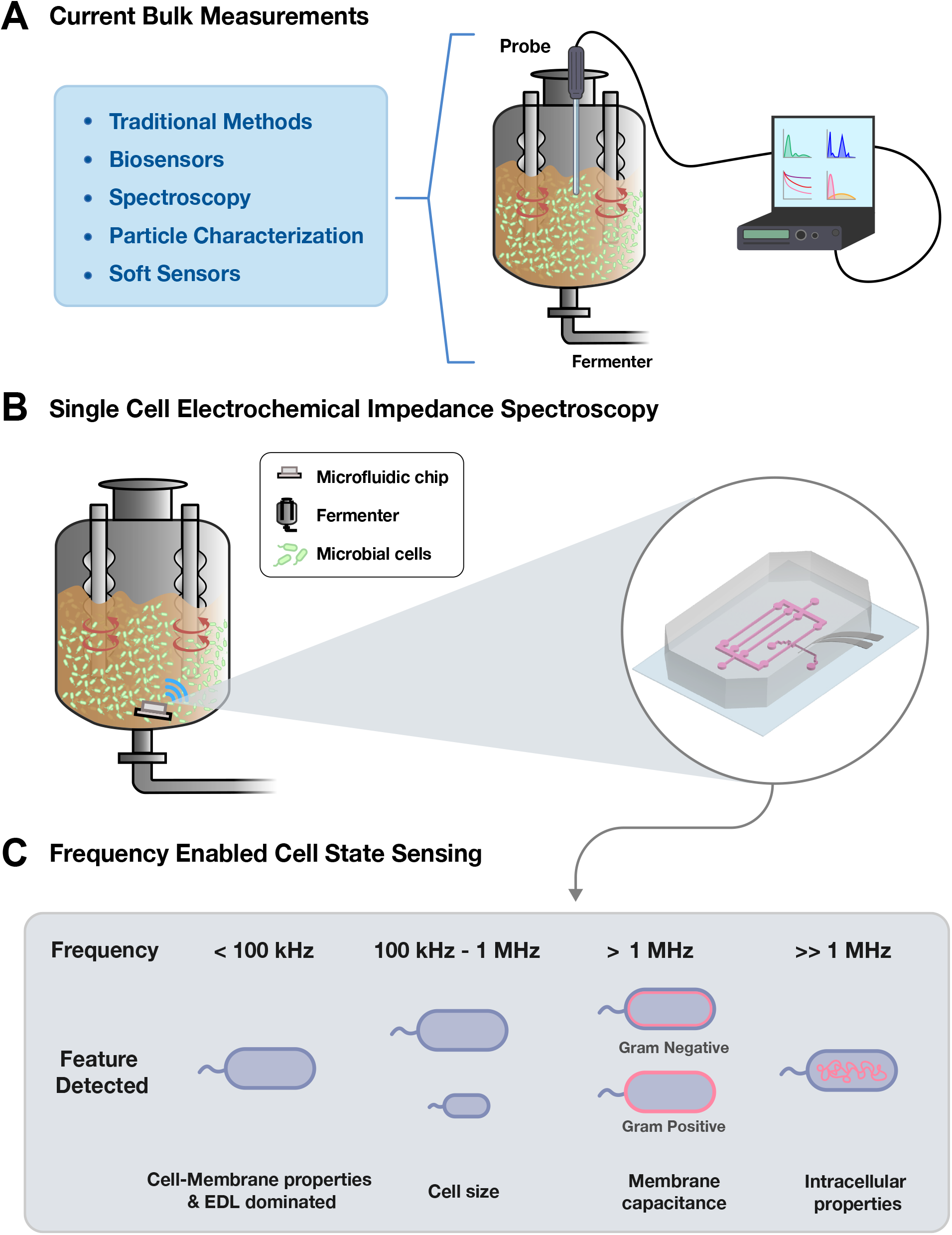
Biomanufacturing concept for single-cell-constrained impedance sensing. **(A)** Different types of current bulk measurements used today include traditional methods (e.g., pH, cell density, dissolved oxygen, and temperature), biosensors, spectroscopy, particle characterization, and soft sensors. **(B)** Future placement of proposed device in-line within the body of the bioreactor. **(C)** Estimated average frequencies used in probing bacterial cells via impedance cytometry, and the corresponding feature being detected.

To address these challenges, we present a microfluidic impedance-sensing platform that directs individual *E. coli*cells from high-density, fermenter-like conditions through a single-cell-constrained sensing path (Figure 1B). This is achieved through passive cell-flux-partitioning geometry coupled with integrated thin-film electrodes for on-chip impedance measurements. In addition to ease of integration within a microfluidic chip, EIS provides the opportunity to probe frequency-dependent electrical properties of biological samples, with different frequency regimes providing sensitivity to features including cell size, membrane capacitance, and intracellular composition (Figure 1C) ^21^. The integration of EIS enables label-free probing of the system while extracting frequency dependent cellular properties. At lower frequencies, the response can reflect cell-membrane polarization, although measurements below several kHz can be dominated by the electrical double layer (EDL) at the electrode-electrolyte interface. At higher frequencies, approximately 100 kHz to 1 MHz provides information related to cell size, frequencies in the few MHz range probe membrane capacitance, and even higher frequencies can provide information on intracellular properties, such as cytoplasmic conductivity ^21^. The design restricts the sensing region to a width comparable to the transverse dimension of an individual *E. coli* cell. To route only a fraction of cells from a high-density input stream toward this sensing region, we developed a passive cell-flux-partitioning architecture implemented as a branched, binary-tree-like network. At successive bifurcations, the incoming flow and associated cell flux are distributed among downstream branches, reducing the fraction of incoming cells routed toward the sensing path without requiring sample preparation or pre-dilution. The platform therefore combines continuous operation with repeated measurements from a restricted sensing path supplied directly from a high-density input culture. Moreover, the proposed chip achieves high reproducibility by using the simplest viable fabrication path compared to more complex alternatives. This platform has the potential to enable reliable, label-free, real-time impedance readouts from a single-cell-constrained flow path operating with high-density microbial cultures.

## 2. Results

### 2.1 Finalized Microfluidic Chip Design

Our target design goal was to develop a system that could take in a high-density cell culture and continuously direct cells through a single-cell-constrained impedance-sensing region without jamming the microfluidic channel. A key design challenge was optimizing the channel geometry to maintain structural stability and reliable cell transport under continuous flow. As no centrifugation or external chip manipulation can be used to position cells inside a fermenter, the flow of the culture is the only driving factor behind any cell movement within the chip. Therefore, the chip requires a channel geometry that passively partitions the incoming cell flux and directs only a fraction of cells toward a single-cell-constrained sensing path. To achieve this, we based our design on a branched network resembling a binary decision tree (Figure 2A). At successive bifurcations, the incoming flow and associated cell flux are distributed among downstream branches, progressively reducing the fraction of cells routed toward the sensing path. A branch then separates from the main channel body and narrows to approximately the transverse dimensions of an individual *E. coli* cell (Figure 2B). Individual cells can therefore be routed toward a sensing region containing electrodes positioned across a cell-constrained flow path while maintaining continuous flow. This region consists of thin-film electrodes deposited on glass and spaced 1 µm apart, enabling integrated impedance measurements from single cell-containing flow through the restricted sensing path (Figure 2C). The branched channel network reduces the fraction of the incoming cell flux directed through the sensing path, enabling individual-cell transit through the electrode region from a high-density input culture. The restricted geometry also increases the relative contribution of cells vs medium within the electrode sensing region compared with a larger unconstrained flow channel.

**Figure 2.**
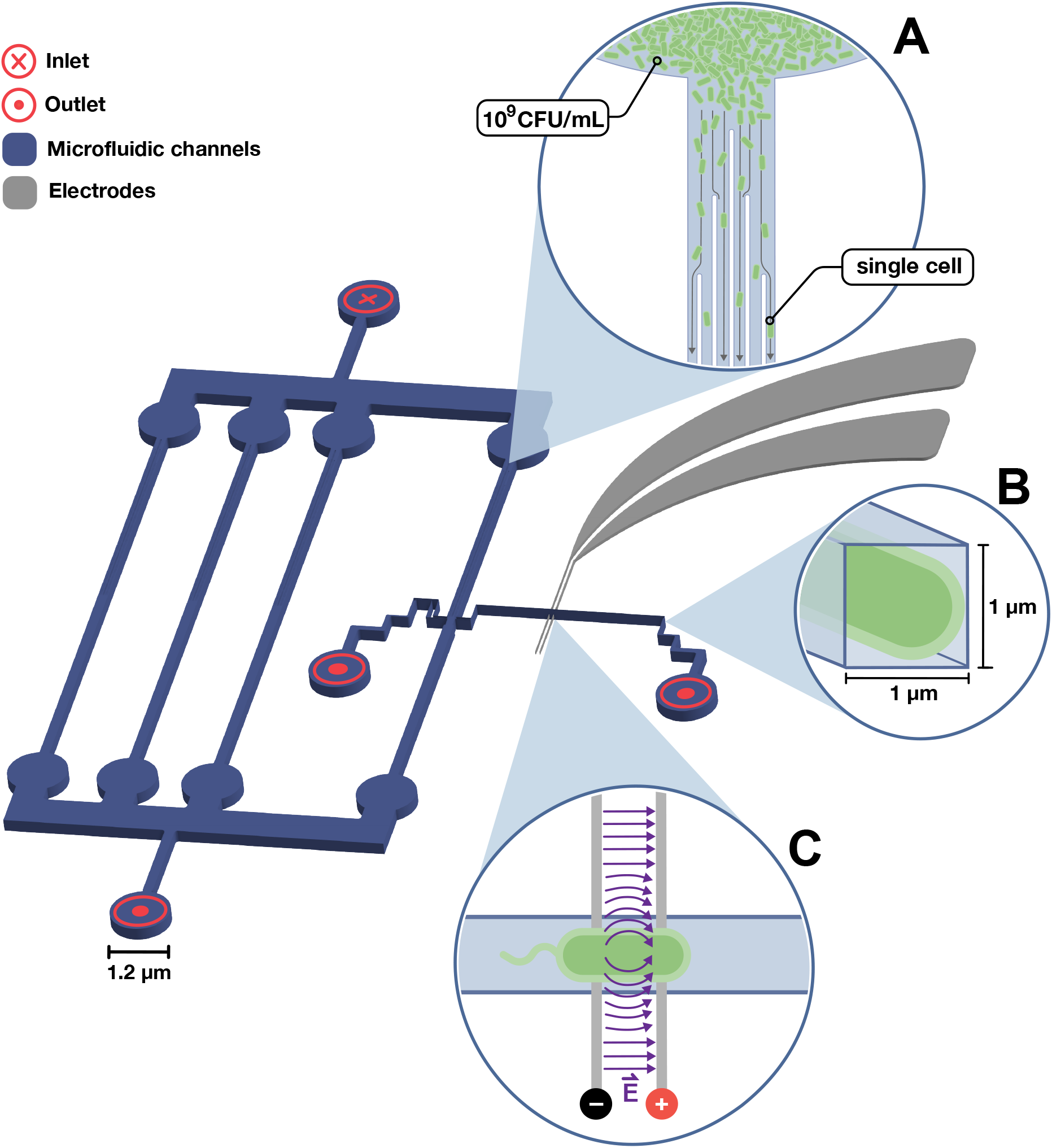
Overview of microfluidic chip and electrode design with key features highlighted. **(A)** Passive cell-flux-partitioning network in which successive channel bifurcations distribute the incoming flow and cell flux across multiple branches, routing a fraction of cells from a high-density input toward the sensing path. **(B)** Measurement channel with a 1 µm × 1 µm cross-section that constrains bacterial transit through the sensing region. **(C)** Integrated impedance-sensing region illustrating a single *E. coli* cell traversing the electric field generated between the two thin-film electrodes.

### 2.2 Chip Fabrication Methodology and Assembly

We fabricated the chips through two parallel work streams that combine traditional photolithography-based PDMS molding and e-beam-based Aluminum electrode patterning on glass (Figure 3A). Following fabrication, the PDMS and glass layers were aligned and bonded using plasma activation and a custom 3D printed alignment jig, achieving permanent sealing verified through leakage testing for up to 72 hours. Profilometer measurements confirmed an average thickness of 0.95 - 1 µm for the resist patterns on the wafer post development, matching design expectations (Figure 3B). Different electrode designs were tested for the deposition of glass, leading to the optimal design consisting of a main testing region between a parallel electrode configuration (Figure 3C). After casting the PDMS and bonding, the fully assembled chip highlights the clear interface between the PDMS microchannels and the electrodes, along with varied electrode positions for testing different regions of the microfluidic chip (Figure 3D). This gives users the flexibility of getting readouts from both single cell areas and bulk population areas if required.

**Figure 3.**
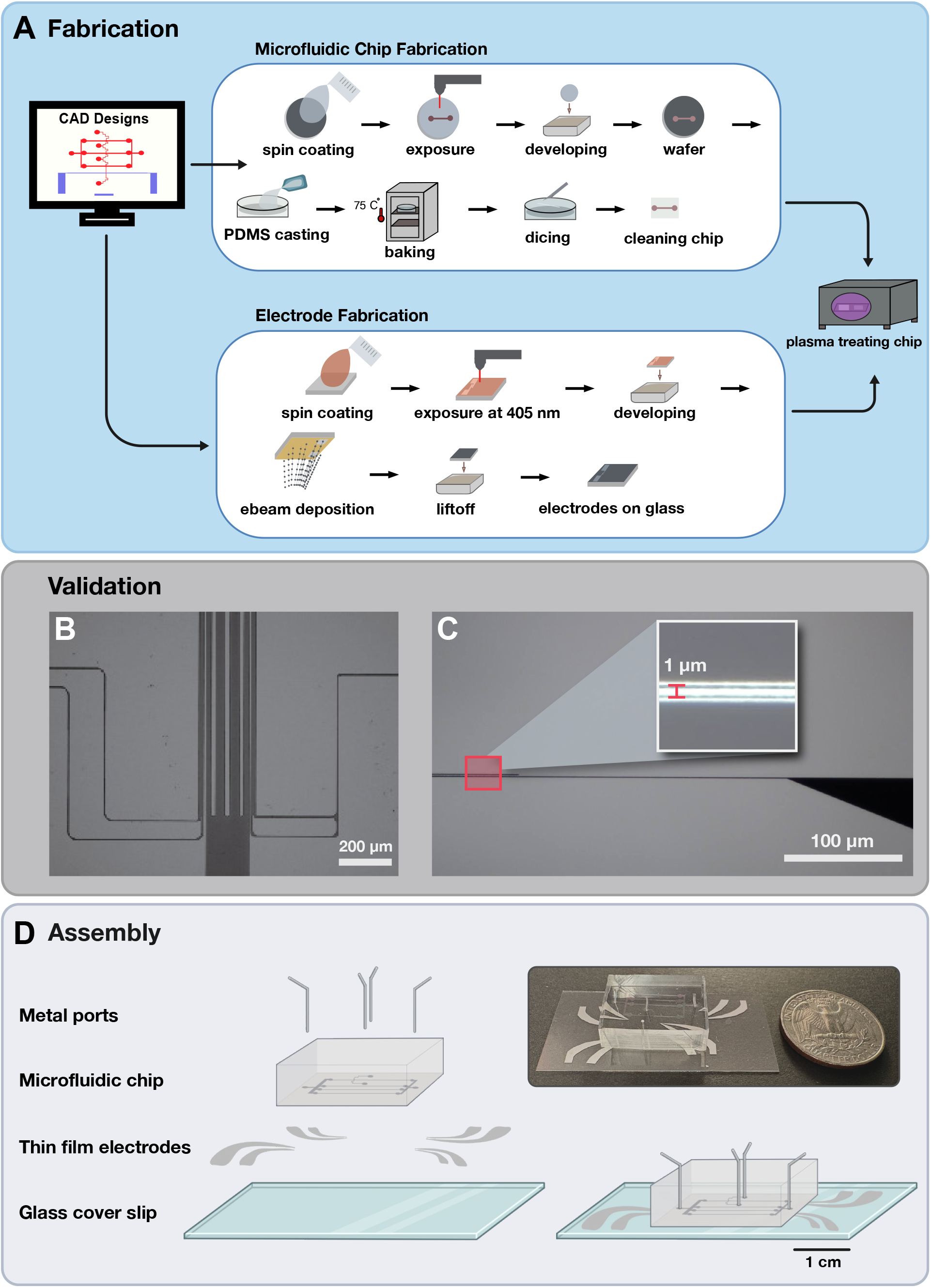
Fabrication and assembly of the integrated microfluidic impedance-sensing device. **(A)** Fabrication roadmap starting from the CAD design of the microfluidic chip, divided into PDMS microchannel fabrication and thin-film electrode deposition on glass substrates, followed by plasma bonding of both parts to each other. **(B)** Microscopic imaging of the photoresist patterns on a Si wafer, confirming successful pattern transfer and dimensional accuracy. **(C)** Verification of electrode deposition on glass, showing uniform thin-film electrodes intact with a 1 µm gap between electrode pairs. **(D)** Final assembled chip illustrating individual layers forming a complete device ready for laboratory testing.

Electrodes placed in the sensing region are particularly sensitive due to the cells’ high-velocity while passing over them. To find the optimal height for the electrodes, multiple iterations were tested in full devices under the same conditions (flow rate, cell type, duration, etc.). The height was optimized to balance electrical sensitivity with hydrodynamic stability. Initial iterations with thicker metal layers ranged from 50-80 nm, which caused flow perturbations and cell blockage effects as cells collided with raised electrodes at the testing region (Figure S3, Movie S2). Conversely, thinner electrodes below 10-15 nm exhibited partial delamination when testing for longer than two hours under constant flow conditions. This resulted in instability or loss of the signal readout. From these results, a thickness of approximately 20 nm was determined to be optimal. Continuous microscopic imaging under operational flow confirmed cell transit through the sensing region without the obstruction observed with earlier electrode configurations for the final device iteration using the 20 nm electrode thickness. This electrode height was used for the validation experiments (see section 2.4). The electrodes generated stable impedance spectra across repeated measurements, validating both their material integrity and electrical stability (Figure S1).

The channel aspect ratio was optimized to minimize PDMS deformation under pressure, ensuring consistency between fabrication batches and reproducibility. Furthermore, the PDMS surface area containing the chip design was selected to ensure stable bonding between the PDMS microfluidic chip and the electrode-patterned glass substrate that seals the device. This resulted in the chip being 2 × 1.3 cm (L x W), while having the bonded and diced PDMS chip be as central as possible on the electrode-patterned glass cover slip. This ensures a more uniform pressure distribution while testing, and therefore reduces the possibility of leakage.

### 2.3 Chip Architecture and Design Iterations

To achieve passive cell-flux partitioning and stable transport through the single-cell-constrained sensing region, several design variants were fabricated and evaluated across two metrics: mechanical robustness of the PDMS microchannels under dynamic pressure gradients (no jamming and/or leakage), and formation of a stable localized electric field within the sensing region for impedance measurements. The COMSOL simulations show the physical channel geometry through multiple iterations leading to the final device (Table 2). All designs progressed in the trend of balancing the geometry with minimum additions of corners that could lead to undesired nonuniform streamlines across different parts of the chip.

**Table 2.**
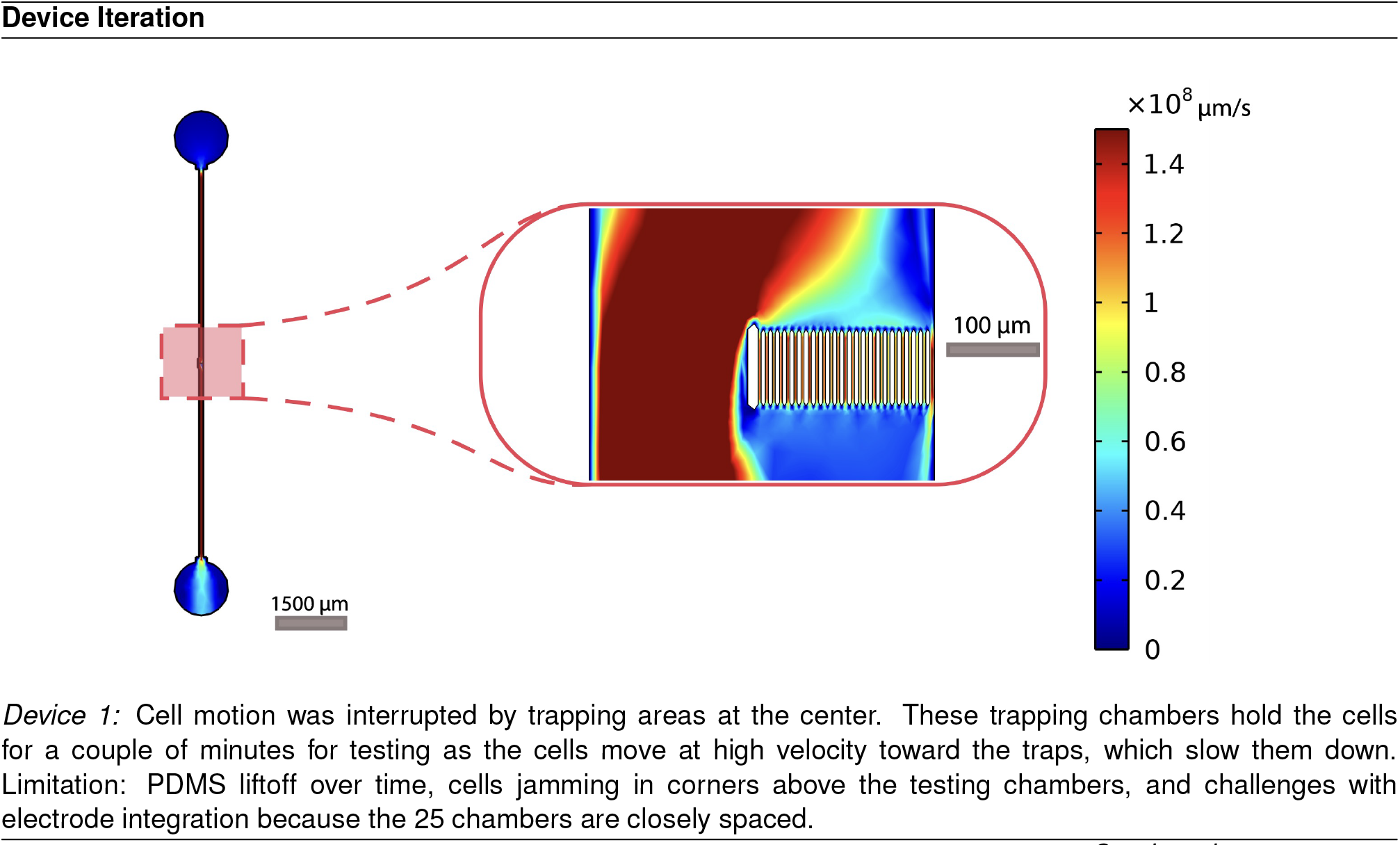
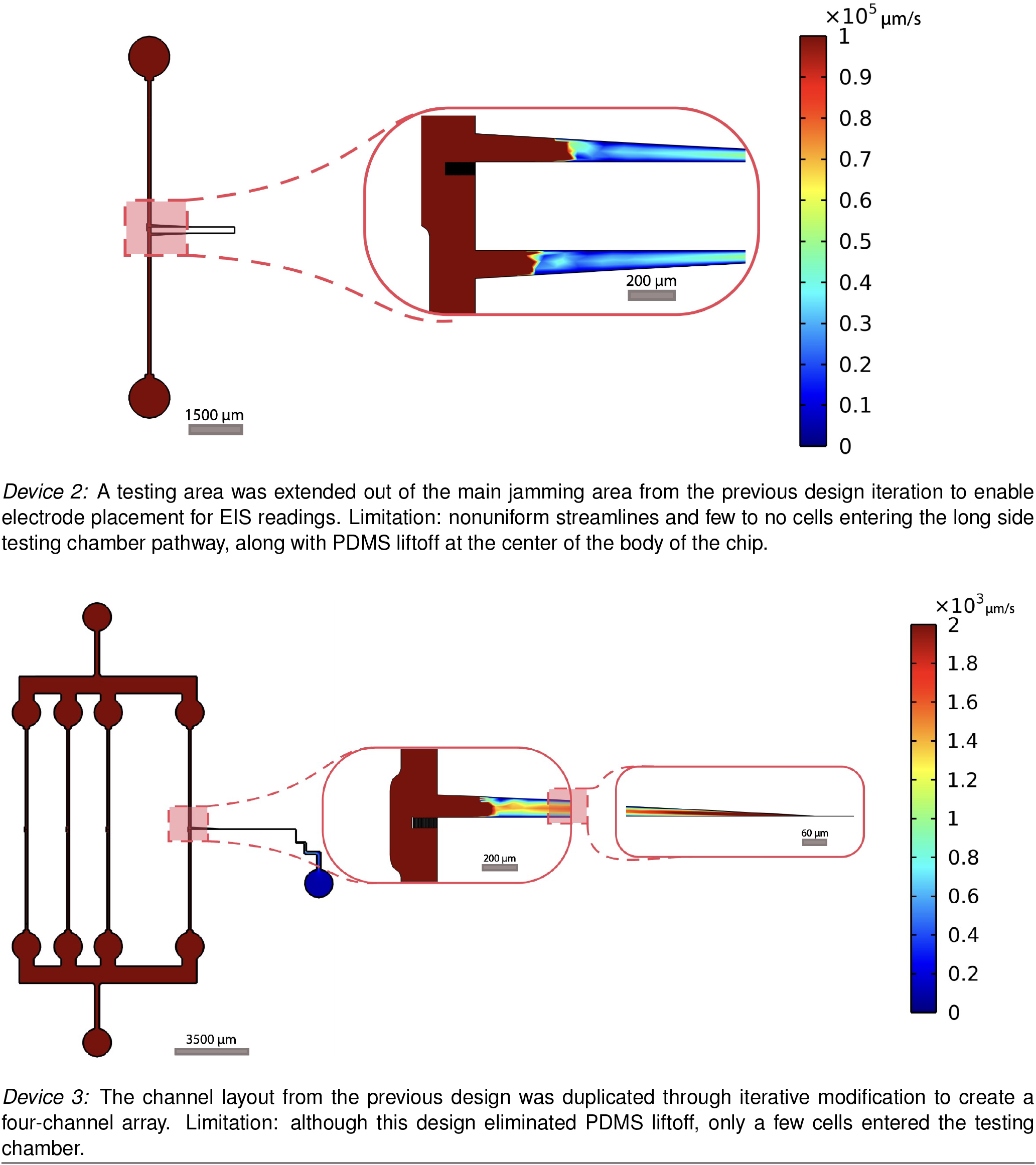
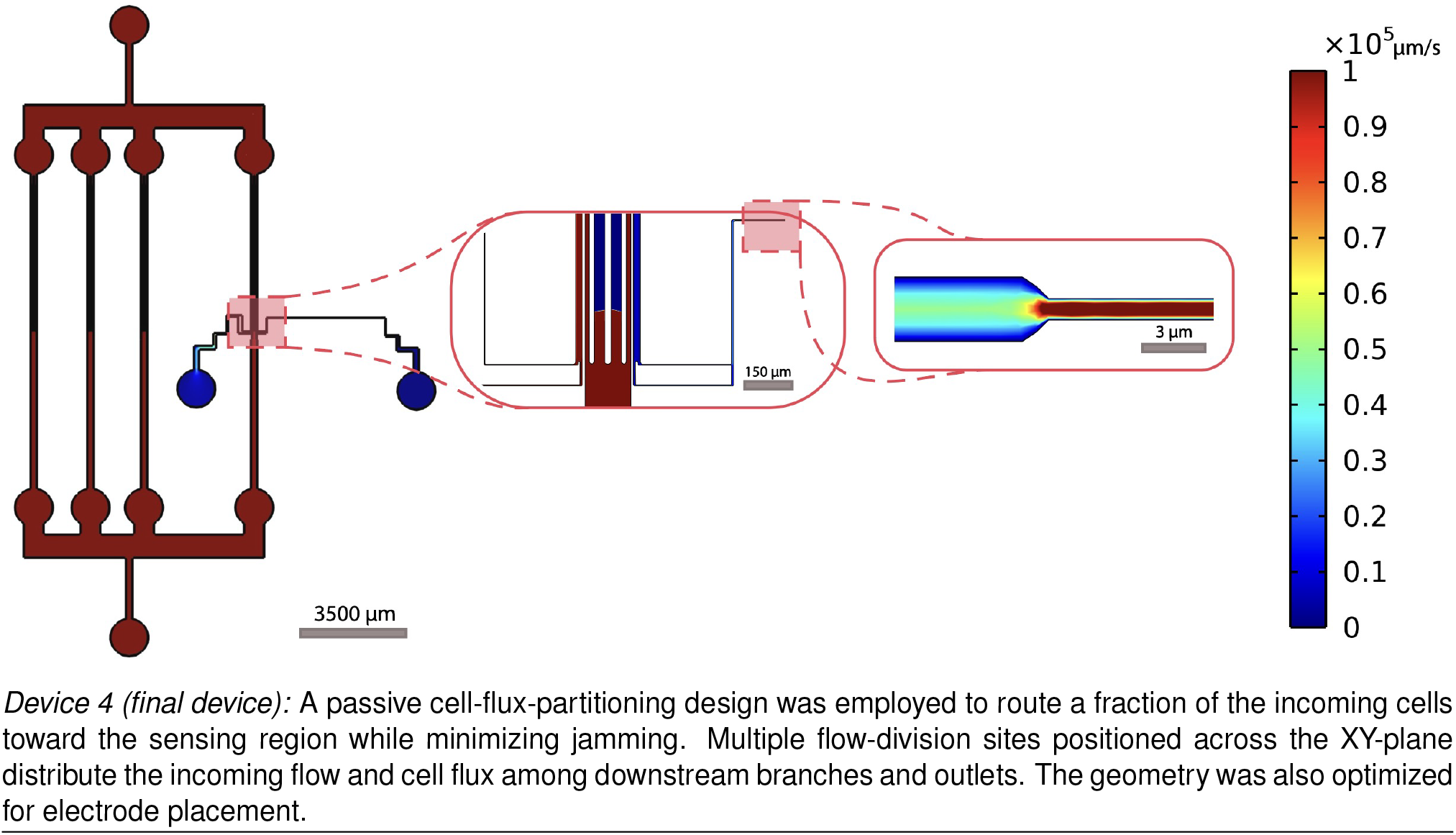
Device iterations with paired testing region simulations for mapping velocity magnitudes.

For Device 1, flow conditions were designed such that only a small portion of cells would be guided into the 25 cell chambers over a short period of time. However, this design exhibited significant shortcomings in practice, primarily in the form of cell jamming above the testing chambers (Figure S4) which prevented reliable operation. To address this, Device 2 introduced a side stream intended to relieve jamming, enable cells to move in a single file, and simultaneously create an additional testing region for electrode placement intersecting this side stream.

This modification enabled cells to move in a single file within a wider region than the 25 central chambers, which could then be used for collective monitoring via adding electrodes to this side stream. However, the geometry of Device 2 produced an unfavorable flow distribution, such that few to no cells entered the intended testing side stream and effective cell-flux partitioning into the sensing branch was not achieved. In addition, the jamming from the first iteration did decrease, but nonuniform flow was observed at the side stream’s inlet. However these two devices together revealed an important biological observation under imaging over time: cells originating from the same culture displayed substantial length variability, even when cell cultures were prepared using identical methods across more than ten individual experimental runs.

Device 3 was developed to address the cells not entering the side stream’s inlet by increasing the cross sectional area of the full chip as a method to mitigate the nonuniform flow. This was accomplished by incorporating duplicated channels to better support the overall chip structure. Across iterations ranging from three to six channels, four channels in the chip body were found to provide the best balance in terms of fabrication feasibility when it comes to the ratio of number of chips to wafer for better turn around time, and leakage performance testing. As the physical PDMS chip’s size was fixed, adding more channels closer to the chip’s sides would cause the chip to detach from the glass coverslip. Nevertheless, even with the revised side stream configuration, Device 3 did not function as intended. The final working configuration, Device 4, redesigned the region preceding the sensing chamber as a branched cell-flux-partitioning network that promotes stable flow into the sensing zone. Successive flow divisions route only a fraction of the incoming cells toward the electrode region, reducing cell traffic through the sensing path while maintaining a high-density input culture. In the final design, the channel divides into asymmetric side branches, one of which contains the sensing region while the other contributes to structural stability and prevents PDMS liftoff. The asymmetric branch geometry produces differential hydraulic resistance that partitions the incoming flow and associated cell flux between the available paths. Through simulation-aided iterative design, this architecture enables passive routing of individual cells from a high-density culture through the restricted sensing region.

### 2.4 Assembled Chip Validation

To evaluate device performance, high-density cultures of *E. coli* at fermenter level concentrations reaching up to 3 × 10^9^ CFU/mL were then run through the chip. The operational parameters summarized in Figure 4B, including the low Reynolds number, are consistent with laminar flow through the device under the tested conditions. EIS measurements were first obtained using LB medium as a background condition, followed by measurements during continuous flow of *E. coli* cell suspensions through the single-cell-constrained sensing path. The spectra in Figure 4C represent the integrated impedance response acquired over an extended measurement period during cell-containing flow. A measurable difference from the background-medium response confirms electrical sensitivity to the presence of cells within the single-cell sensing path. To illustrate the basis of the cell-flux-partitioning behavior, Figure 4D overlays a hydraulic-resistance network on the chip geometry, showing the relative fluidic resistances across the channel segments (R_3_ > R_1_ > R_5_ > R_2_ > R_4_). These resistance differences govern the distribution of flow, and therefore the fraction of incoming cell flux directed through each downstream branch. Additionally, the overall fluidics behavior of the system was governed by laminar flow dynamics consistent with Hagen-Poiseuille’s law. These design elements provide a stable, restricted flow path through the electrode sensing region. Figure 4E and the corresponding Movie S1 independently demonstrates the corresponding microfluidic behavior, with a representative fluorescent *E. coli* cell imaged traversing the parallel-electrode region under continuous flow. This optical observation demonstrates passive routing of individual cells through the sensing path from a high-density input using only geometric channel features.

**Figure 4.**
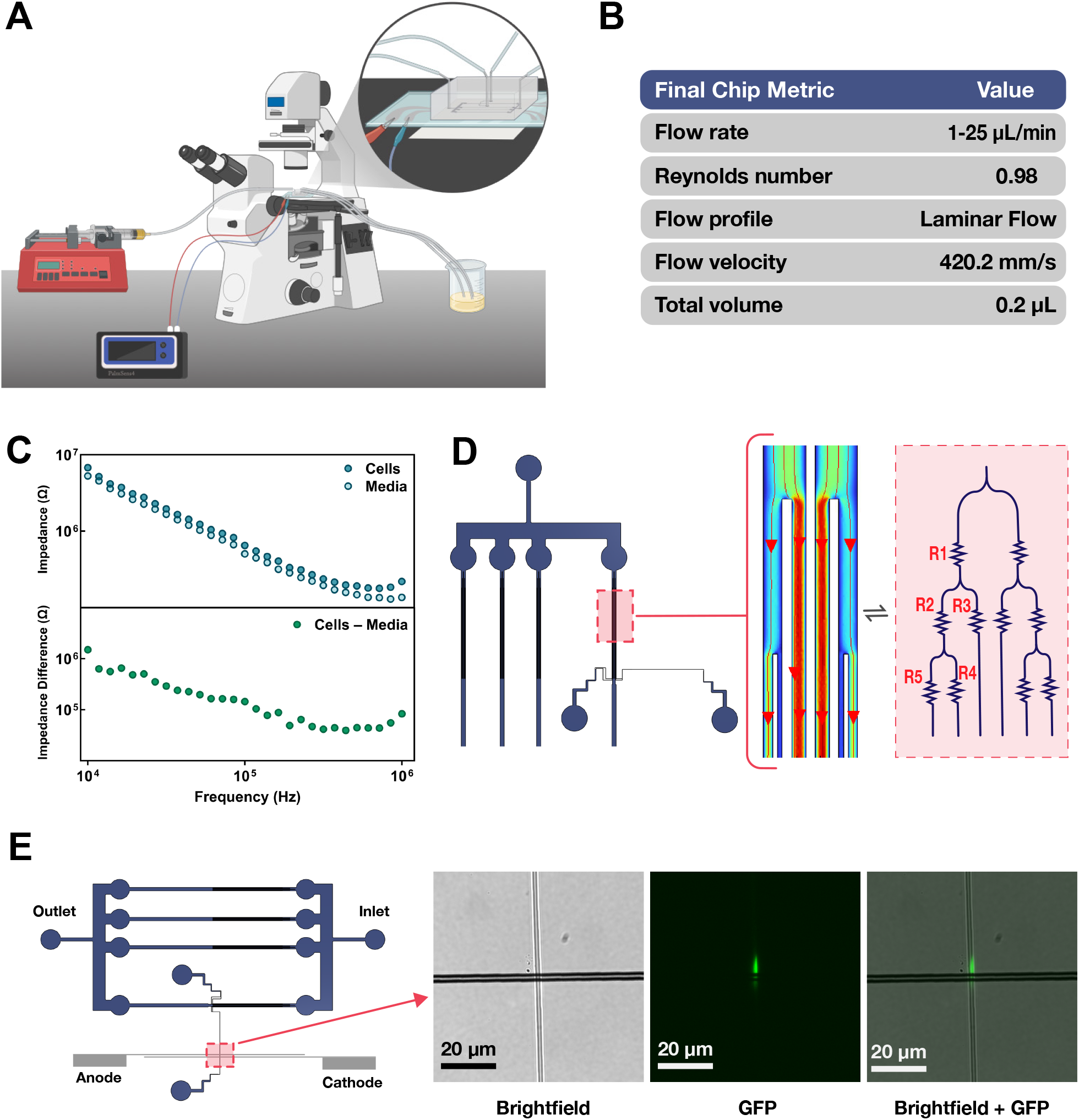
Experimental validation of cell-flux partitioning and integrated impedance sensing. **(A)** Full experimental setup showing syringe pumps connected to the microfluidic chip and the PalmSens4 device attached to the selected electrode pair for testing within the 1 µm chamber. Three outlets are connected to a waste container. **(B)** Operational flow parameters for the final chip configuration. **(C)** The time-averaged impedance response acquired during continuous flow of *E. coli* through the single-cell-constrained sensing path and the corresponding LB-medium background response, as well as the impedance difference between the two conditions. **(D)** COMSOL flow lines of the channel structure with an fluidic circuit analogy overlaid, showing the relationship R_3_ > R_1_ > R_5_ > R_2_ > R_4_. **(E)** Representative image of a fluorescent *E. coli* cell traversing the electrode sensing region under brightfield, GFP, and merged imaging.

## 3. Discussion

A central challenge in translating single-cell analytical technologies to microbial biomanufacturing is the mismatch between the high cell densities required for productive fermentation and the sparse, spatially constrained cell flow required for measurements at cellular resolution. Biomanufacturing is increasingly important for the commercial production of food and other valuable products ^22^, while scale-up places a growing premium on continuous monitoring strategies capable of providing information about microbial populations during production ^23,24^. Here, we address this mismatch with a passive microfluidic architecture that partitions cell flux from a high-density input culture before directing cells through a single-cell-constrained impedance-sensing region. The device operates continuously with microbial inputs reaching 10^9^ cells/mL without requiring upstream dilution or active cell sorting. Integrated thin-film electrodes distinguish cell-containing flow from the corresponding medium background, while fluorescence microscopy independently demonstrates individual-cell transit through the restricted sensing region. Together, these results establish a microfluidic interface between high-density microbial cultures and a sensing geometry defined at the scale of individual cells.

The key advance of this architecture is control over the cell flux presented to the analytical region. This is important for bioprocess monitoring because the upstream culture can remain at process-relevant density while the branched microfluidic network establishes a downstream transport regime compatible with a single-cell-scale sensing geometry. Single-cell analytical approaches can then reveal cellular properties and population heterogeneity that are obscured in bulk measurements ^11,25^. In the present device, differential hydraulic resistance across successive branches distributes the incoming flow and associated cell flux among multiple paths, routing only a fraction of incoming cells toward the sensing region. Because this partitioning arises from channel geometry alone, it avoids centrifugation, externally actuated sorting, or other active sample-handling steps. Although demonstrated here using integrated impedance electrodes, the underlying cell-flux-partitioning strategy could provide a general microfluidic front end for other microscale analytical modalities whose performance is limited by the need for sample processing to achieve compatible cell fluxes.

Established impedance cytometry approaches can use transient electrical signals to characterize cellular properties ^26^, but these measurements generally rely on sufficiently separated cell transits to associate electrical perturbations with individual cells. At high cell concentrations, simultaneous or closely spaced cell passages can compromise event-level interpretation and create additional challenges associated with transport and channel obstruction ^27–29^. The present implementation addresses the upstream fluidic component of this problem by restricting the cell traffic delivered to the sensing region rather than diluting the original culture. Impedance spectra in this study are integrated over the acquisition period and therefore characterize cell-containing flow through the constrained sensing path rather than individual cell-transit events. The parallel thin-film electrode configuration however provides a defined sensing region of about 1 × 1 × 3 µm (WxHxL) through which cells pass ^30^, and prior impedance-cytometry studies demonstrate that, with appropriate calibration and event-resolved acquisition, electrical measurements can report parameters including cell size, membrane capacitance, and cytoplasmic conductivity ^26,31–33^. The present platform therefore establishes the fluidic and device-level interface needed to bring such measurements into contact with high-density microbial feeds.

Several steps remain before this architecture can function as a fully integrated bioprocess analytical technology. Future work should determine whether the local hydrodynamic conditions experienced during passage through the restricted sensing region measurably alter microbial physiological state, particularly for measurements intended to report native cell properties. On the electrical side, synchronized optical and impedance acquisition, higher-bandwidth measurements, and calibration against defined cellular states will be required to associate impedance perturbations with individual cells. Longer-term deployment may additionally benefit from alternative electrode materials with greater stability, wireless or remotely integrated readout electronics, and fluidic strategies that support direct coupling to bioreactor sampling streams.

By separating process-scale cell density from the cell traffic presented to a microscale analytical region, our passive cell-flux partitioning strategy provides a route for connecting high-density microbial bioprocesses with sensing technologies that operate most effectively at cellular length scales. This creates a path toward biomanufacturing measurements that move beyond bulk environmental variables toward direct interrogation of microbial populations during production.

## 4. Methods

### 4.1 Materials

All photoresists, developers, and diluents mentioned were obtained from Kayaku Advanced Materials, Inc. and Thermo Fisher Scientific. Four-inch silicon wafers were purchased from University Wafer. Microscope coverslips were Fisherbrand cover glasses with dimensions of 50 24 0.17 mm. Restriction enzymes, HiFi DNA Assembly Master Mix (NEB #E2621S), and chemically competent *Escherichia coli* cells (NEB #C2987I) were obtained from New England Biolabs (NEB; Ipswich, MA, USA). Growth-media components were obtained from BD Biosciences (Franklin Lakes, NJ, USA), and synthetic DNA constructs were purchased from Integrated DNA Technologies (IDT; Coralville, IA, USA).

### 4.2 Plasmid Cloning

The fluorescent *E. coli* strain used in this work was made by transforming plasmid pGL003-pVeg1-GFP(sp) into*E. coli* strain C2987 (NEB). The plasmid pGL003-pVeg1-GFP(sp) was generated by inserting the constitutive *Bacillus subtilis* promoter pVeg1 upstream of a GFP sequence codon-optimized for *Streptococcus pneumoniae* in the parent plasmid pGL003. While this construct is designed for *B. subtilis*, we found that it generates a very strong signal in *E. coli*, and therefore we use it for *E. coli* tracking in this work. The vector pGL003 was a gift from Gene-Wei Li (Addgene plasmid #215335). Plasmids were cloned and amplified in *E. coli* strain C2987 (NEB).

*E. coli* transformations, HiFi DNA assembly reactions, gel extractions, and minipreps were performed using kits from New England Biolabs (NEB; Ipswich, MA, USA). All other commercial chemical reagents were obtained from Sigma-Aldrich (St. Louis, MO, USA). Standard isothermal DNA assembly was performed using the NEB HiFi DNA Assembly Kit. A double-stranded DNA fragment (IDT) and the pGL003 parent plasmid (Addgene #215335), prepared using standard methods, were used as input DNA. Sequence-verified plasmids (Plasmidsaurus) were transformed into chemically competent *E. coli* strain C2987 (NEB #C2987I) according to the manufacturer’s recommendations. Protocols were based on the product instructions for the NEB HiFi DNA Assembly Master Mix (NEB #E2621S) and *E. coli* chemically competent cells (NEB #C2987I). The full plasmid sequence of pGL003-pVeg1-GFP(sp) is provided in the Supplementary Material.

### 4.3 Cell Culturing

Routine biomass generation was performed by culturing cells in liquid LB medium at 37 °C with shaking at 270 rpm. When indicated, LB medium used to culture plasmid-bearing *E. coli* was supplemented with ampicillin at 100 µg mL^−1^. To mimic fermenter-scale cell concentrations, cultures were grown to stationary phase for 18 h, reaching an approximate concentration of 3 × 10^9^ CFU/mL. For optical validation of cell transit through the electrode sensing region, fluorescent *E. coli* cultures were grown for 12 h to an approximate concentration of 3 × 10^8^ CFU/mL. For leakage, durability, and jamming experiments, stationary-phase cultures were concentrated 35-fold by centrifugation to generate maximal-density cultures. Optical-density measurements at 600 nm (OD_600_) and fluorescence measurements were acquired using an Infinite M200 plate reader (Tecan).

### 4.4 Chip Fabrication

#### Wafer Patterning

CAD drawings were created in KLayout to define the network of microchannels: boundaries, inlet/outlet ports, and testing chambers. GDSII files were used in an MLA150 maskless aligner, by placing a 4-inch Silicon wafer to pattern the designs onto. This wafer is pretreated with oxygen plasma for cleaning the wafer using 300 mL/min O2, 300 W for 5 mins (Asher, PVA TePla America M4L). Then a spin coated layer is placed on the wafer (PWM32-PS-CB15 Spinner,Headway Research, Inc.) using photoresist SU-8 2002 6:4 in cyclopentanone mixture. Spin coating steps are as follows: (a) 500 RPM for 10 seconds, (b) 5000 RPM for 30 seconds, (c) deceleration to 0 RPM over 2 seconds. Then, the wafer is soft-baked on a hotplate at 65 °C for 30 seconds and 95 °C for 3 minutes before exposure. Next, expose the resist in the MLA150 maskless aligner (Heidelberg Instruments), using 1950 mJ/cm^2^ exposure with a defocus of -6 via the 375 nm writing head. Post exposure, baking was performed on a hotplate at 65 °C for 1.5 minutes and then 95 °C for 2 minutes. Followed by development, by placing the wafer in a bath of SU8 developer for 30 seconds and then washed with Isopropanol then Acetone each for 3 seconds maximum, then dried with N2 gas.

#### Soft Lithography And Chip Dicing

PDMS (10:1 base:curing agent) was mixed and degassed under vacuum for 30 minutes (Krayden Dow Sylgard 184 Silicone Elastometer Kit. Clear, Fisher Scientific), poured over the fabricated wafer, cured by baking at 75 °C for 12 hours, peeled off by dicing, and ports were punched with a 0.75 mm inner diameter biopsy puncher (Electron Microscopy Sciences). Then, appropriate metal ports (Instech Stainless steel catheter coupler, 22ga x 15mm, non-sterile) were inserted, along with appropriate tubing (Accuflow Systems TYG TUB.020 ID 500FT-150MPK Fisher Scientific 50-236-5803) to prepare for connecting the chip with a syringe pump.

#### Electrodes on glass deposition

The fabrication process described here represents the primary and most used approach in this study. Using a maskless aligner (MLA150), the CAD designed electrode patterns were transferred onto a photoresist mold in preparation for subsequent electron-beam metal deposition. The procedure was conducted as follows. Glass coverslips were first cleaned using oxygen plasma at a flow rate of 300 mL/min O2 and a power of 300 W for 5 minutes (Asher, PVA TePla America M4L). Immediately following plasma cleaning, three drops of Bis(trimethylsilyl)amine (HMDS) were dispensed onto the center of each coverslip to promote adhesion between the glass surface and the photoresist layer during the subsequent coating step. Photoresist S1813 was spin-coated at 4000 RPM, followed by a pre-exposure bake at 115 °C for 4 minutes. Samples were then exposed using a 405 nm writing head with an exposure dose of 180 mj/cm^2^ and a defocus setting of -4. Development was performed via a bath of MF319 developer for 40 seconds, after which the samples were baked again on the same hotplate for 3 minutes to complete post-exposure hardening. The prepared substrates were subsequently placed in the electron-beam deposition system (Electron Beam and Thermal Evaporator, CHA Solutions) (Figure S2). Aluminum was deposited at a controlled rate between 1 Å/s and 5 Å/s to achieve thin-film layers of approximately 20 nm thickness. Following metal deposition, lift-off was performed using sonication at 40 kHz (Branson Ultrasonic Cleaners, Model 1800) in an acetone bath (Acetone LM) for 1-2 minutes. The completed electrode substrates were then rinsed, dried, and prepared for bonding to the PDMS microchannel layer to form the final integrated device.

### 4.5 Bonding

Diced PDMS chips and glass coverslips with patterned electrodes were bonded using a plasma bonder (Harrick PDC-001 plasma cleaner), at the low setting level for 20 seconds. After plasma treatment, a 3D-printed alignment jig was used to position the microfluidic features and glass cover slip with the patterned electrodes exactly perpendicular to the channel they bisected, to avoid misalignment and ensure reproducibility.

### 4.6 COMSOL Simulations

Computational validation of the microfluidic design was performed using the steady-state conditions of a 2D laminar flow model implemented in COMSOL Multiphysics. The computational model employed was a depth-averaged single-phase laminar flow formulation. Device geometries were imported from CAD layouts and repaired prior to simulation. The microchannel height was set to 1 µm, and fluid properties corresponding to water were used for density and viscosity. A constant volumetric flow rate of 25 µL/min was applied at the inlet, while all remaining channel terminations were treated as pressure outlets with zero gauge pressure. No-slip boundary conditions were applied at all channel walls, and incompressible flow was assumed. Simulations were performed using a stationary study with a refined mesh and a direct PARDISO solver to ensure numerical stability. Flow behavior was evaluated using streamline visualizations and pressure distribution maps along channel walls to assess flow uniformity and pressure gradients within the sensing region.

### 4.7 Experimental Setup and EIS Measurements

Prior to each experiment, the microfluidic chip was primed by infusing media at a higher flow rate to ensure complete filling of the channels and removal of trapped air, after which the flow rate was reduced to 1.5–3 µL/min for the duration of the experiment. Experiments were conducted for approximately 48-72 hours on average. The chip was positioned on an inverted microscope (Leica DMi8), and GFP fluorescence was used to confirm the presence of cells and verify that cells were passing through the testing region. EIS measurements were acquired using a PalmSens4 potentiostat connected to the chip electrodes in the appropriate working electrode (WE), reference electrode (RE), and counter electrode (CE) configuration. The PalmSens4 measurement file was configured to acquire impedance magnitude (Z) as a function of time. Prior to each experiment, the measurement frequency and total acquisition duration were adjusted as needed for the specific experimental conditions, while the remaining measurement parameters were kept consistent. Impedance spectra were acquired over a frequency range of 1 kHz to 1 MHz using 43 logarithmically spaced frequency points (14 points per decade) on average. Measurements were performed with a DC bias of 0 V and a sinusoidal AC amplitude of 0.25 V RMS. Each frequency sweep required approximately 37 s to complete. Three consecutive EIS sweeps were performed for each measurement, and the impedance values obtained from the three sweeps were averaged at each frequency for subsequent analysis.

## Supporting information

pGL003-pVeg1-GFP(sp).gb

## Additional Information

### DATA AVAILABILITY

All data needed to evaluate conclusions in the paper are present in the paper and/or the supplementary information.

### AUTHOR CONTRIBUTIONS

*Author contributions according to the CRediT taxonomy are as follows:*

**W.A**.: Conceptualization, Data curation, Formal analysis, Investigation, Methodology, Project administration, Validation, Visualization, Writing—original draft, and Writing—review & editing.

**L.P**.: Conceptualization, Formal analysis, Methodology, Visualization, Writing—original draft, and Writing—review & editing.

**Y.P**.: Data curation, Formal analysis, and Writing—review & editing.

**J.J.R.Y**.: Validation.

**Y.Y**.: Project administration, Resources, and Writing—review & editing.

**R.Y**.: Resources, Supervision, and Writing—review & editing.

**M.J**.: Conceptualization, Funding acquisition, Investigation, Methodology, Project administration, Resources, Supervision, Validation, Visualization, and Writing—review & editing.

All authors read and approved the final manuscript.

## ACKNOWLEDGEMENTS

The authors acknowledge the use of the Boston University Photonics Center shared facilities. This work was performed in part using the shared laboratories and instrumentation supported by the Boston University Photonics Center.

W.A. was funded by Aramco through the College Preparatory Program (CPP).

## References

[1] Tom A. Ewing, Niels Nouse, Matthijs van Lint, Jacco van Haveren, Jeroen Hugenholtz, and Daan S. van Es. Fermentation for the production of biobased chemicals in a circular economy: a perspective for the period 2022-2050. Green Chemistry, 24(17):6373–6405, 2022. ISSN 1463-9262. doi: 10.1039/d1gc04758b.

[2] Nelson Mota de Carvalho, Eduardo M. Costa, Sara Silva, Lígia Pimentel, Tito H. Fernandes, and Manuela Estevez Pintado. Fermented foods and beverages in human diet and their influence on gut microbiota and health. Fermentation, 4(4), 2018. ISSN 2311-5637. doi: 10.3390/fermentation4040090.

[3] Mallari Praveen and Simone Brogi. Microbial Fermentation in Food and Beverage Industries: Innovations, Challenges, and Opportunities. Foods, 14(1):114, January 2025. ISSN 2304-8158. doi: 10.3390/foods14010114.

[4] R. Takors. Scale-up of microbial processes: Impacts, tools and open questions. Journal of Biotechnology, 160(1):3–9, July 2012. ISSN 0168-1656. doi: 10.1016/j.jbiotec.2011.12.010.

[5] Frank Delvigne, Ralf Takors, Rob Mudde, Walter van Gulik, and Henk Noorman. Bioprocess scale-up/down as integrative enabling technology: From fluid mechanics to systems biology and beyond. Microbial Biotechnology, 10(5):1267–1274, 2017. ISSN 1751-7915. doi: 10.1111/1751-7915.12803.

[6] Jianye Xia, Guan Wang, Jihan Lin, Yonghong Wang, Ju Chu, Yingping Zhuang, and Siliang Zhang. Advances and practices of bioprocess scale-up. In Jie Bao, Qin Ye, and Jian-Jiang Zhong, editors, Bioreactor Engineering Research and Industrial Applications II, pages 137–151. Springer Berlin Heidelberg, Berlin, Heidelberg, 2016. ISBN 978-3-662-48347-3. doi: 10.1007/10_2014_293.

[7] Jianye Xia, Guan Wang, Meng Fan, Min Chen, Zeyu Wang, and Yingping Zhuang. Understanding the scale-up of fermentation processes from the viewpoint of the flow field in bioreactors and the physiological response of strains. Chinese Journal of Chemical Engineering, 30:178–184, February 2021. ISSN 1004-9541. doi: 10.1016/j.cjche.2020.12.004.

[8] Carina L Gargalo, Isuru Udugama, Katrin Pontius, Pau C Lopez, Rasmus F Nielsen, Aliyeh Hasanzadeh, Seyed Soheil Mansouri, Christoph Bayer, Helena Junicke, and Krist V Gernaey. Towards smart biomanufacturing: A perspective on recent developments in industrial measurement and monitoring technologies for bio-based production processes. Journal of Industrial Microbiology and Biotechnology, 47(11):947–964, November 2020. ISSN 1367-5435. doi: 10.1007/s10295-020-02308-1.

[9] Peter Harms, Yordan Kostov, and Govind Rao. Bioprocess monitoring. Current Opinion in Biotechnology, 13(2):124–127, 2002. ISSN 0958-1669. doi: 10.1016/S0958-1669(02)00295-1.

[10] Hu Zhang, Xiaolin Cui, Jingxiu Bi, Sheng Dai, and Haitao Ye. Single-cell analysis for bioprocessing. Engineering in Life Sciences, 15(6):582–592, 2015. ISSN 1618-2863. doi: 10.1002/elsc.201400155.

[11] Yang He, Yuehui He, Yuanyuan Zhou, Xunrong Li, Xinran Zhang, Yuetong Ji, Junhong Yu, and Jian Xu. Tracking production and interconversion of extra- and intra-cellular metabolites during beer fermentation by ramanomics. Bioresource Technology, 443:133788, March 2026. ISSN 0960-8524. doi: 10.1016/j.biortech.2025.133788.

[12] Xi Sun, Xin Zhou, Ran Yu, Xiaofang Zhou, Jun Zhang, Teng Xu, Jianmei Wang, Mengqi Li, Xiaoting Li, Min Zhang, Jian Xu, and Jia Zhang. Assessing the physiological properties of baker’s yeast based on single-cell Raman spectrum technology. Synthetic and Systems Biotechnology, 10(1):110–118, March 2025. ISSN 2405-805X. doi: 10.1016/j.synbio.2024.09.004.

[13] Nicole A. Dambruin, Jack T. Pronk, and Marieke E. Klijn. Application of process analytical technology for real-time monitoring of synthetic co-culture bioprocesses. Analytical and Bioanalytical Chemistry, 417(25):5611–5625, June 2025. ISSN 1618-2650. doi: 10.1007/s00216-025-05949-2.

[14] Felipe Tapia, Daniel Vázquez-Ramírez, Yvonne Genzel, and Udo Reichl. Bioreactors for high cell density and continuous multi-stage cultivations: Options for process intensification in cell culture-based viral vaccine production. Applied Microbiology and Biotechnology, 100:2121–2132, 2016. ISSN 0175-7598. doi: 10.1007/s00253-015-7267-9.

[15] Anne Y. Fu, Charles Spence, Axel Scherer, Frances H. Arnold, and Stephen R. Quake. A microfabricated fluorescence-activated cell sorter. Nature Biotechnology, 17(11):1109–1111, November 1999. ISSN 1546-1696. doi: 10.1038/15095.

[16] Sharath Narayana Iyengar, Tharagan Kumar, Gustaf Mårtensson, and Aman Russom. High resolution and rapid separation of bacteria from blood using elasto-inertial microfluidics. ELECTROPHORESIS, 42(23):2538–2551, 2021. doi: 10.1002/elps.202100140.

[17] L. D’Amico, N. J. Ajami, J. A. Adachi, P. R. C. Gascoyne, and J. F. Petrosino. Isolation and concentration of bacteria from blood using microfluidic membraneless dialysis and dielectrophoresis. Lab on a Chip, 17(7):1340–1348, April 2017. ISSN 1473-0197. doi: 10.1039/c6lc01277a.

[18] Avinash V Police Patil, Yu-Sheng Chuang, Che-Wei Lin, Chiou-Ying Yang, Tomoyuki Yasukawa, and Ching-Chou Wu. A Microfluidic Platform Integrating Dielectrophoretic Concentration and Impedimetric Sensing for Rapid Salmonella choleraesuis Detection. ACS Sensors, 10(9):6512–6520, September 2025. doi: 10.1021/acssensors.5c00779.

[19] Jiahong Chen, Jianwei Zhong, Hongtao Lei, and Ye Ai. Label-free multidimensional bacterial characterization with an ultrawide detectable concentration range by microfluidic impedance cytometry. Lab on a Chip, 23(23):5029–5038, December 2023. ISSN 1473-0197. doi: 10.1039/d3lc00799e.

[20] Wenshuai Wu, Gaozhe Cai, Yang Liu, Yuanjie Suo, Boran Zhang, Wei Jin, Yinghua Yu, and Ying Mu. Direct single-cell antimicrobial susceptibility testing of escherichia coli in urine using a ready-to-use 3d microwell array chip. Lab on a Chip, 23(10):2399–2410, 05 2023. ISSN 1473-0197. doi: 10.1039/d2lc01095j.

[21] Zhao Zhang, Xiaowen Huang, Ke Liu, Tiancong Lan, Zixin Wang, and Zhen Zhu. Recent Advances in Electrical Impedance Sensing Technology for Single-Cell Analysis. Biosensors, 11(11):470, November 2021. ISSN 2079-6374. doi: 10.3390/bios11110470.

[22] Shahida Anusha Siddiqui, Zeki Erol, Jerina Rugji, Fulya Taşçi, Hatice Ahu Kahraman, Valeria Toppi, Laura Musa, Giacomo Di Giacinto, Nur Alim Bahmid, Mohammad Mehdizadeh, and Roberto Castro-Muñoz. An overview of fermentation in the food industry - looking back from a new perspective. Bioresources and Bioprocessing, 10(1):85, November 2023. ISSN 2197-4365. doi: 10.1186/s40643-023-00702-y.

[23] Roger Boulton, James Nelson, and André Knoesen. A Review of Sensors for the Monitoring, Modeling, and Control of Commercial Wine Fermentations. Fermentation, 11(6):329, June 2025. ISSN 2311-5637. doi: 10.3390/fermentation11060329.

[24] Joan Cortada-Garcia, Jennifer Haggarty, Stefan Weidt, Rónán Daly, S. Alison Arnold, and Karl Burgess. On-line targeted metabolomics for real-time monitoring of relevant compounds in fermentation processes. Biotechnology and Bioengineering, 121(2):683–695, February 2024. ISSN 0006-3592. doi: 10.1002/bit.28599.

[25] Byron F. Brehm-Stecher and Eric A. Johnson. Single-Cell Microbiology: Tools, Technologies, and Applications. Microbiology and Molecular Biology Reviews, 68(3):538–559, September 2004. ISSN 1092-2172. doi: 10.1128/MMBR.68.3.538-559.2004.

[26] Chayakorn Petchakup, King Ho Holden Li, and Han Wei Hou. Advances in Single Cell Impedance Cytometry for Biomedical Applications. Micromachines, 8(3):87, March 2017. ISSN 2072-666X. doi: 10.3390/mi8030087.

[27] Carlos Honrado, Paolo Bisegna, Nathan S. Swami, and Federica Caselli. Single-cell microfluidic impedance cytometry: from raw signals to cell phenotypes using data analytics. Lab on a Chip, 21(1):22–54, 01 2021. ISSN 1473-0197. doi: 10.1039/d0lc00840k.

[28] Yuwen Li, Can Huang, Song-I. Han, and Arum Han. Measurement of dielectric properties of cells at single-cell resolution using electrorotation. Biomedical Microdevices, 24(2):23, June 2022. ISSN 1572-8781. doi: 10.1007/s10544-022-00621-3.

[29] Shu Zhu, Xiaozhe Zhang, Zheng Zhou, Yu Han, Nan Xiang, and Zhonghua Ni. Microfluidic impedance cytometry for single-cell sensing: Review on electrode configurations. Talanta, 233:122571, October 2021. ISSN 0039-9140. doi: 10.1016/j.talanta.2021.122571.

[30] Mahtab Kokabi, Gulam M. Rather, and Mehdi Javanmard. Ionic Cell Microscopy: A new modality for visualizing cells using microfluidic impedance cytometry and generative artificial intelligence. Biosensors and Bioelectronics: X, 24:100619, August 2025. ISSN 2590-1370. doi: 10.1016/j.biosx.2025.100619.

[31] Yongxiang Feng, Zhen Cheng, Huichao Chai, Weihua He, Liang Huang, and Wenhui Wang. Neural network-enhanced real-time impedance flow cytometry for single-cell intrinsic characterization. Lab on a Chip, 22(2):240–249, 01 2022. ISSN 1473-0197. doi: 10.1039/d1lc00755f.

[32] Quentin Rezard, Faruk Azam Shaik, Jean Claude Gerbedoen, Fabrizio Cleri, Dominique Collard, Chann Lagadec, and Mehmet C. Tarhan. Silicon Compatible Process To Integrate Impedance Cytometry With Mechanical Characterization. In 2023 IEEE 36th International Conference on Micro Electro Mechanical Systems (MEMS), pages 475–478, January 2023. doi: 10.1109/MEMS49605.2023.10052621.

[33] Fenni Zhang, Shaopeng Wang, Yunze Yang, Jiapei Jiang, and Nongjian Tao. Imaging single bacterial cells with electro-optical impedance microscopy. ACS sensors, 6(2):348–354, February 2021. ISSN 2379-3694. doi: 10.1021/acssensors.0c00751.

