## Supplementary Material for "Passive Cell-Flux Partitioning Enables Single-Cell-Constrained Impedance Sensing From High-Density Microbial Cultures"

### The PDF file includes:

- fig. S1. Electrode performance comparison.
- fig. S2. Electrode deposition setup.
- fig. S3. Cell passage through the testing region.
- fig. S4. Cell jamming.

### Other Supplementary Material for this manuscript includes the following:

- movie S1 (.mp4 format). Cell movement from the main branch through the testing region.
- movie S2 (.mp4 format). Cell displacement from the electrode surface due to increased electrode thickness.
- pGL003-pVeg1-GFP(sp) (.gb format). Plasmid map for GFP expression in *E. coli*.

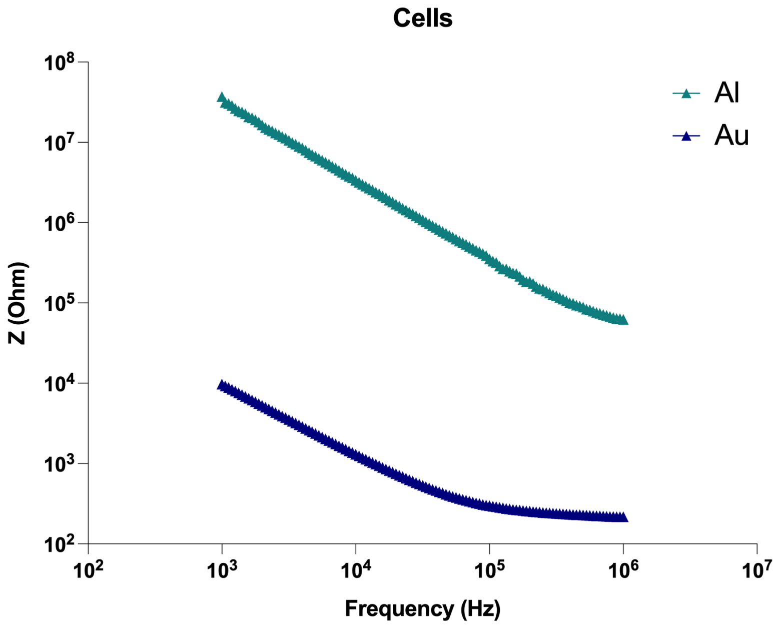


**Figure S1:** Impedance response comparison between aluminum parallel electrodes fabricated in this device and commercially available gold interdigitated electrodes. Measurements were performed using samples from the same cell culture to compare EIS signal trends. The comparable response patterns indicate structural integrity of the fabricated electrodes, while differences in signal magnitude are attributed to the distinct electrical resistivities of the electrode materials and differing electrode geometries.

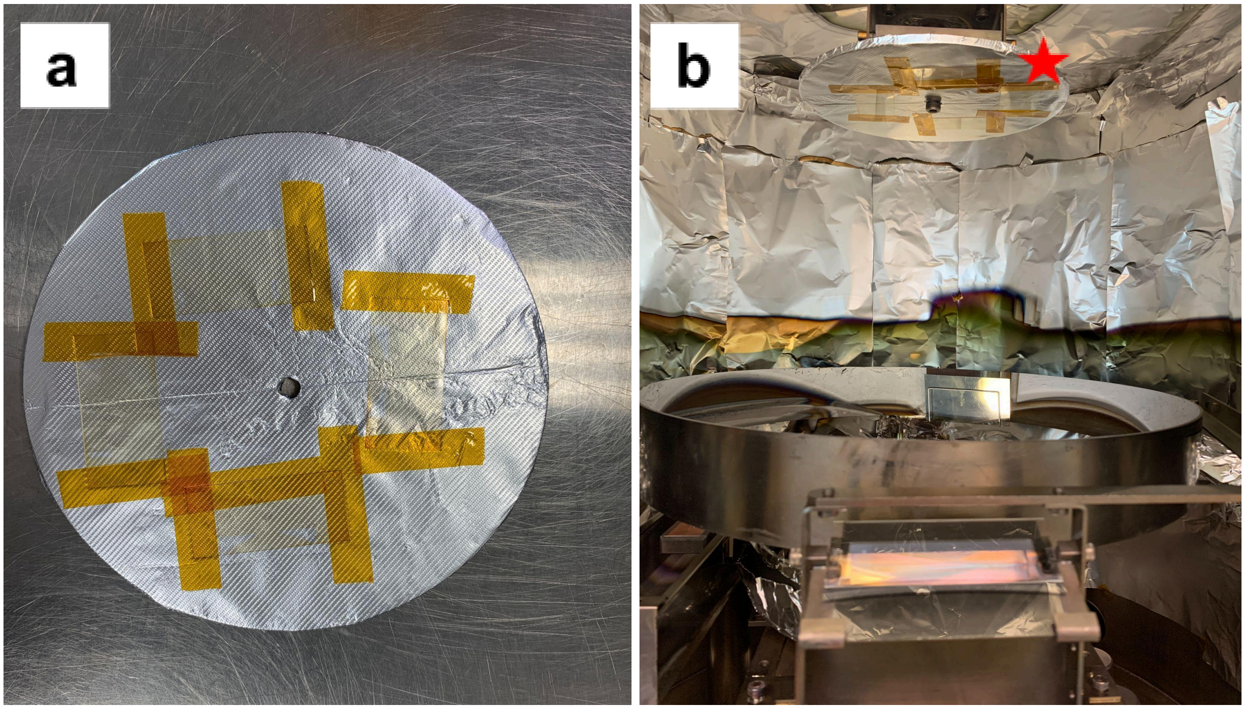


**Figure S2:** (a) Positioning of the patterned glass coverslips on the holding plate following photolithography for deposition of thin-film aluminum electrodes. (b) Placement of the glass coverslip holding plate inside the e-beam deposition system, indicated by the red star.

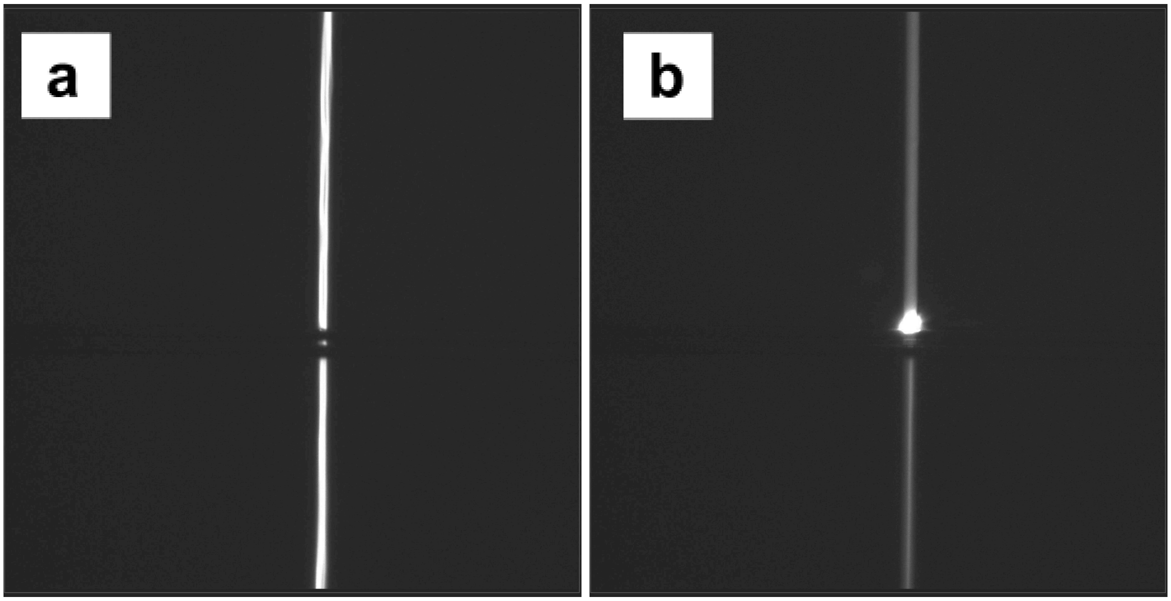


**Figure S3:** Effect of increased electrode thickness on cell transit through the sensing region. (a,b) Representative GFP images acquired with 50–80 nm-thick electrodes show cells approaching and interacting with the raised electrode region. The increased electrode height perturbs cell transit (b) relative to the final 20-nm configuration, including displacement of cells near the electrode surface. The corresponding dynamic behavior is shown in Movie S2.

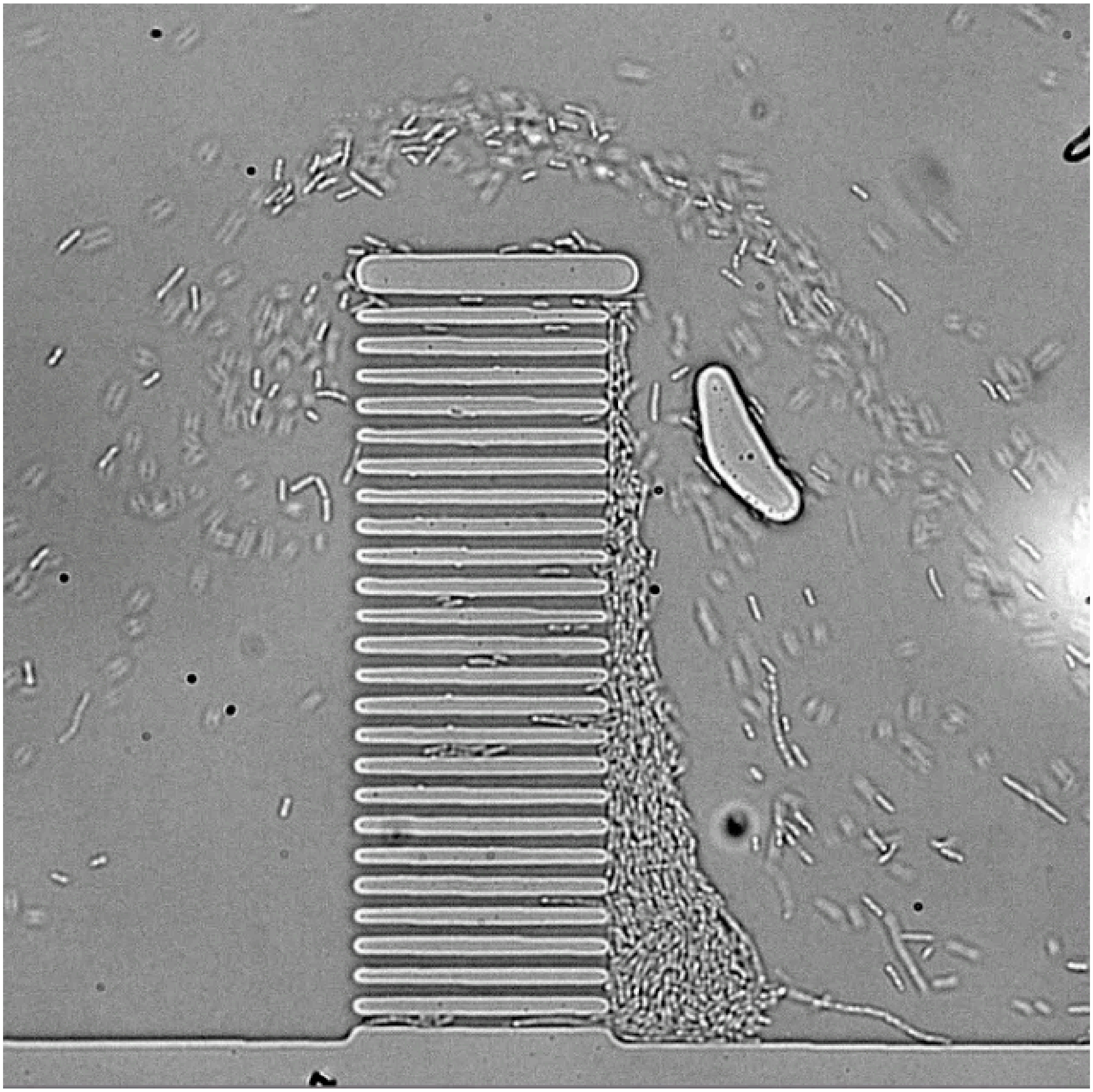

**Figure S4:** Cell jamming in Device 1. Cells initially enter the central trapping/sensing region, but persistent accumulation develops around the chamber array during continuous flow. The resulting jamming and nonuniform cell distribution prevented reliable operation and motivated redesign of the channel architecture.

**Supplementary file captions:**

- **movie S1. Cell movement through the testing region.** Cells moving from the main branch of the microfluidic channel and passing through the electrode EIS based testing region during continuous flow.

- **movie S2. Cell displacement from thick electrodes.** Cells approaching the testing region and being displaced away from the electrode surface as they encounter electrodes with increased thickness.

- **pGL003-pVeg1-GFP(sp) (.gb).** GenBank file containing the complete plasmid sequence used for constitutive GFP expression in this study.
